# Observation of Field Cancerization in Human Breast Tissue with Raman Hyperspectral Imaging and Artificial Neural Networks

**DOI:** 10.1101/2025.03.23.643829

**Authors:** J. Nicholas Taylor, Kazuki Bando, Yasuto Naoi, Satoshi Fujita

## Abstract

Tissue-conserving partial mastectomy surgeries are often used to treat breast cancer, in which malignancies and surrounding margin tissues are removed. Morphologies of margin tissues are often assessed intraoperatively by a histopathologist, requiring excision and time-consuming preparation for microscopic examination. Changes in morphology often appear only after the mutation of cells involved in tumor propagation, suggesting that margins and perhaps conserved tissue may already contain cancer-converted cells prior to morphological changes. In a process known as field cancerization (FC), benign tissue surrounding a malignancy is affected by continued exposure to carcinogens, triggering cancer-promoting alterations like conversions of normal adipocytes and fibroblasts to cancer-associated adipocytes and fibroblasts. We report successful FC detection in human breast tissue samples excised during breast-conserving surgeries in human patients using Raman hyperspectral imaging and artificial neural networks (ANNs). Spectra in the images are segmented based on tissue modalities such as adipose, connective, and glandular tissues using intensities at Raman shifts of known molecules and label propagation, a method of transductive learning. ANNs that discriminate cancer from noncancer spectra return high quality (>95% accuracy) and reproducible distinction of spectra originating from cancerous tissue at high signal-to-error ratios (S/Es). Good distinction is returned in all modalities, with adipose-rich modalities returning the most reliable classifications, suggesting reliable discrimination of normal and cancer-associated adipocytes. Spectral S/Es are increased using larger spectral measurement areas, but this causes spectral mixing across different tissue modalities and significant biological variation, failing to deliver a corresponding increase in classification quality.

## INTRODUCTION

Treatment of cancerous tumors with conservation of breast tissue involves surgical excision of the malignant tissue as well as histopathological assessment and possible excision of surrounding tissues^1^. Tissue surrounding a tumor is often excised and assessed intraoperatively for malignancy by a qualified histopathologist, requiring visual recognition of morphological changes in the tissue sections that are associated with oncogenic transformation^2–4^. In various types of breast cancer, malignancies can occur through oncogenic transformation of luminal epithelial (LE) cells in human breast tissue (HBT), which often undergo large-scale epigenetic and phenotypic changes in a process known as the endothelial-mesenchymal transition (EMT)^5–7^. LEs can change phenotypes to basal-like myoepithelial (ME) cells, acquiring new capabilities that allow mutated cells to anchor to the basement membrane of the extracellular matrix (ECM) and act as stem cells for tumor propagation after oncogenesis ^8,9^. Although the EMT alters LE cellular morphologies and allows for histopathological recognition, non-epithelial tissue components surrounding the malignancy are also altered in an effect known as field cancerization (FC) ^10,11^. FC is a phenomenon in which the cells and tissue surrounding malignant regions are altered by continued exposure to carcinogens associated with cancer development^12^. FC can result in widespread epigenetic and molecular alterations to normal HBT, altering gene expressions through processes such as DNA methylation and transforming the nearby cells and tissues to cancer-associated cells and tissues^13^.

HBT contains several distinct tissue modalities; stromal modalities provide structural and metabolic support for glandular modalities in which milk is stored and produced^14–17^. Glandular tissue modalities are interconnected networks of tubular ducts that bifurcate to form branched structures called lobules, clusters of 10-100 rounded sacs called acini where milk is stored and produced. LEs line the interiors of the ducts and acini, and are encased by MEs, which anchor the glandular modality to the stroma and contract during lactation to expel milk. The stromal modality contains the fibrous support structure of the ECM as well as adipose tissue and fibroblasts, and other modalities like blood vessels and immune cells such as macrophages and lymphocytes. The ECM, composed primarily of fibrous collagen and other structural proteins such as elastin and fibronectin, is subdivided into the interstitial matrix and the basement membrane, which anchors to the MEs of the glandular modality. Dispersed throughout the ECM are fibroblasts, cells that are responsible for production of ECM components, as well as participation in tissue development, homeostasis, and immune regulation^18^. Finally, adipocytes are primary components of the adipose modality, fat-storing cells that serve several functions in HBT, including structural and metabolic support of glandular modalities, releasing adipokines and other growth factors that regulate endocrine functions in HBT^19^.

Widespread epigenetic changes due to FC lead to a host of morphological and molecular changes in cancerous HBT ^11^. In glandular modalities, although MEs are less affected by FC than other cell types, the number of MEs in cancerous tissue has been observed to decrease relative to normal HBT as cancer progresses^20,21^, resulting in weaker barriers against invading cells and LEs that are more susceptible to oncogenesis. FC also causes alterations outside the glandular tissues^22^. Stiffness of the ECM increases through changes in composition, with differing ratios of collagen, fibronectin, and elastin^23^, and increased crosslinking among structures^24^. Fibroblasts can transform into cancer-associated fibroblasts (CAFs) that promote cancer progression^25,26^ through the secretion of growth factors and cytokines, enhancement of angiogenesis, induction of the epithelial-mesenchymal transition, and induction of oxidative stress^27^. Adipocytes can transform to cancer-associated adipocytes (CAAs) that also promote cancer growth^28–30^, having irregular shapes with smaller, dispersed lipid droplets, remodeling the ECM through changes in shapes. CAAs also provide metabolic support^31^ and growth factors that enhance metastasis and promote the endothelial-mesenchymal transition in LEs^32^.

Raman spectroscopy has shown recognition of FC^33^ and Raman hyperspectral imaging (RHI) is known to provide non-destructive, non-invasive characterizations of microscopic structures and properties across many types of systems^34^, including applications from food science^35,36^ to mineralogy^37^. Applications to both *in vitro* and *in vivo* biological systems are extensive, with Raman spectroscopic applications of stimulated^38^ and coherent anti-Stokes^39^ Raman scattering having been developed for use in microscopic imaging of biological systems. RHI using spontaneous Raman scattering^34^ has been used to characterize biological properties of non-alcoholic fatty liver disease^40,41^ and thyroid cancer cell lines^42^, for example, as well as demonstrating a bevy of uses in characterization of breast cancer^43^. RHI has often been used in combination with machine learning (ML)^44^, characterizing measurements for background contributions^45^, increasing measurement speed^46^, standardizing measurements across devices, and for discrimination of various disease and tissue types^47,48^. Recently, artificial neural networks (ANNs) have been implemented to characterize Raman spectra^44^, e.g., those that are trained monitor tumor responses to radiation therapy^49^ along with other ANN approaches that seek to discriminate, for example, spectra originating from bodily fluids of various species^50^.

Raman hyperspectral imaging is used in combination with ANNs and other ML approaches to characterize normal and malignant HBT that was excised during surgeries to treat occurrences of breast cancer. Label propagation, a method of transductive ML in which partial information is propagated to estimate properties of unlabeled data^51^, is used to segment spectra into collagen-rich, adipose-rich, and non-stromal groups that approximately represent primary HBT modalities such as the ECM, adipocytes, and glandular tissues, respectively. To analyze and assess the suitability of RHI for use in the characterization of HBT and the discrimination of normal and malignant HBT, ANNs are built and trained to classify Raman hyperspectra as cancer or noncancer. Posterior evaluation of ANN model outputs and ground truth histopathological assignments show reasonable classification accuracies and good recovery of spectral properties of cancer/noncancer (C/NC) tissues. ANN model outputs are observed to be dependent on HBT modality, as spectra originating from adipose-rich tissues return the highest C/NC classification qualities (CQs). This result suggests accurate distinction normal adipocytes from cancer-associated adipocytes (CAAs) and the successful of detection of field cancerization with RHI and ANNs. Further examination of CQ versus signal-to-error (S/E) ratios indicate C/NC classification approaching perfect accuracy in all HBT modalities at high S/E. Higher S/E spectra are obtained from adipose-rich tissues, owing to large Raman scattering cross sections, which is the source of higher CQ in the adipose-rich segment. In general, these results suggest robust detection of malignancies and/or cancer field effects in all HBT modalities, given that Raman spectra are of sufficiently high quality as quantified by S/E.

## METHODS

### Properties of HBT Modalities in Histological and Raman Hyperspectral Images

Raman hyperspectral images are used to gain information about local chemical microenvironments. Raman spectra are useful because they contain contributions from many relevant biomolecules, such as proteins, lipids, and DNA^52^. Microscopic imaging gives insight into spatial relationships among the molecules and provides visualization of the morphological structures of the sample. Local prominences of important molecules can be inferred through their relative Raman scattering intensities, allowing for identification of various tissue modalities, such as adipose-rich tissues through spectral bands associated with hydrocarbon chains. Local conditions can also be relayed through changes in intensities of molecules that are susceptible to changes in those conditions. For example, carotenoids are susceptible to oxidation and will show reduced Raman intensities when under oxidative stress.

Fig. 1 compares a histological image to several Raman images of HBT. Fig. 1A shows a representative hematoxylin and eosin (H&E) stained histological image of HBT, with collagen-rich stroma, adipocytes, and lobular structures indicated within, while Fig. 1B shows 4 composite images of HBT obtained using Raman microscopy. First, we note that there is a vast difference in physical scale between the histological and Raman images, as the observation area in the histological image is approximately 1,000× larger than the areas of the Raman images. Whereas H&E staining represents adipocytes (white), collagen-rich stroma (pink), glandular regions (purple), and morphological features with ease, Fig. 1B relays considerable chemical detail in the Raman hyperspectral images (RHIs). Four prominent molecular types are displayed in the RHIs of Fig. 1B using 4 color scales that correspond to normalized Raman intensities in the indicated ranges: cytochromes (pyrrole breathing, 740-760 cm^-1^, blue), collagen (amide III stretching, 1240-1260 cm^-1^, pink), carotenoids (polyene vibration, 1510-1530 cm^-1^, orange), and lipids (CH_2_ symmetric stretching, 2840- 2860 cm^-1^, yellow). Black color represents low intensity while brighter colors represent larger intensities. Colors at each pixel are mixed as intensity-weighted averages of the component colors in a standard RGB color space. A total of 319 RHIs are included in this work, acquired from normal and cancerous HBT samples that were excised during breast cancer surgeries of 40 individual patients. The 319 RHIs contain 957,000 Raman hyperspectra, each of which represents a 1.5×1.5 *μ*m^2^ region of excised HBT. Cancerous (red) and noncancerous (blue) averaged Raman spectra, over all patients and RHIs, are shown in Fig. 1C, along with spectral positions and representative spatial mappings of each of the 4 molecular components.

**Figure 1.**
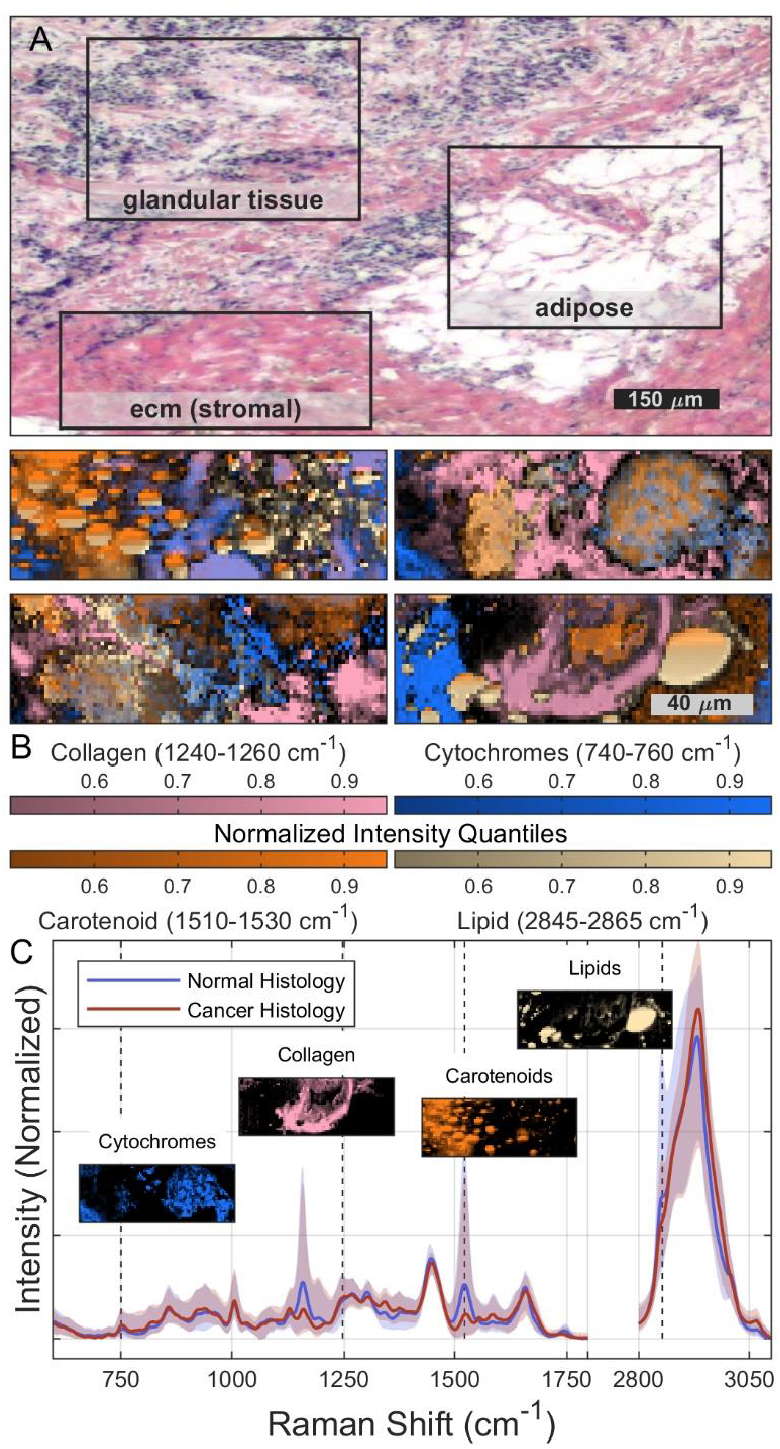
Human breast tissue (HBT) modalities. A) A H&E-stained microscopic image used for histopathological assessments indicating regions of the ECM (pink), adipose (white), and glandular (purple) tissue modalities. B) Intensity-weighted mixtures of Raman intensities observed from cytochromes (740-760 cm^-1^, blue), collagen (1240-1260 cm^-1^, pink), carotenoids (1510-1530 cm^-1^, orange), and lipids (2845-2865 cm^-1^, yellow), are shown in the spatial dimensions of 4 Raman images. Component intensities are mapped to the color scales shown in the legend and mixed in an RGB-color space according to their normalized spectral intensities. C) Spectral overlay showing average spectra (solid lines) and their IQRs (shaded regions) for samples assigned cancer (red) and normal (blue) histology by a histopathologist. Representative spatial mappings of the intensities of important molecules are included near their spectral locations. Intensities are conveyed in appropriate colors in the legend of B).

### Label Propagation for Recognition of HBT Modalities

Although each sample in this study is histologically labeled as cancer or noncancer by a pathologist, pointwise labels of HBT modalities within the RHIs are not available. Rather, Raman intensities of prominent molecules in HBT modalities are used to generate sets of initial labels for label propagation (LP), a transductive machine learning (ML) method that propagates a partial set of labels throughout the unlabeled members of the data set. LP is an iterative classifier in which labels are propagated using, in this case, a *k*-nearest-neighbors model in a principal component space, as detailed in the Appendix and in supporting Fig. S1. Spectra are segmented with LP for recognition of high collagen intensities, indicating collagen-rich tissue modalities that most likely originate in the ECM, as well as for recognition of adipose-rich modalities. 96 full LP repetitions (reps) were performed for recognition of collagen-rich and adipose-rich spectra, such that each spectrum is labeled as either positive or negative for collagen and either positive or negative for adipose. Little overlap occurs among positive collagen-rich and adipose-rich segments over the 96 reps; an average of only 497 of 957,000 total spectra were positive for both collagen-rich and adipose-rich segments. For these spectra, segmentation labels are adjusted randomly in post-processing such that each spectrum carries either a positive collagen label or a positive adipose label, but not both. A third segment arises from LP segmentation: spectra that return negative for both collagen-rich and adipose-rich segmentation. These spectra, termed the non-stromal segment, are regarded as primarily containing glandular tissue modalities, although other components such as blood vessels and immune cells may occupy this segment as well. We note that some flexibility in spectral segment interpretations is necessary in the absence of a ground truth segmentation of HBT modalities.

Spatial mappings of selected LP segmentations are shown in Fig. 2A, indicating contiguous regions of collagen-rich (pink), adipose-rich (yellow), and non-stromal (purple) segments, suggesting that LP sufficiently partitions similar spectra from the targeted HBT modalities. Collagen-rich (upper panel), adipose-rich (center panel), and non-stromal (lower panel) average spectra (solid lines) and their interquartile ranges (IQRs, shaded regions) over 96 LP reps are shown in Fig. 2B, with collagen-rich and adipose-rich spectra showing characteristics common to fibrous collagen spectra and lipid spectra, respectively. Finally, Fig. 2C shows a ternary plot of the populations of each segment within each RHI. Small circles, colored red for cancerous samples and blue for non-cancerous, represent segment populations of individual RHIs in individual LP reps, while larger red and blue markers indicate the average segment population of each RHI over all LP reps. Green circular markers represent the average segment populations aggregated over all RHIs within each of the 96 LP reps. As shown in Fig. 2C, while average segment populations (green markers) show little variation over the LP reps, populations in individual images vary widely, indicating that the spatial sizes of the RHIs are typically not large enough to recover the average relative populations of HBT modalities. It is also of note that little population bias is observed between cancerous and noncancerous samples, as dispersion of the populations of each tissue type are relatively consistent across the population space.

**Figure 2.**
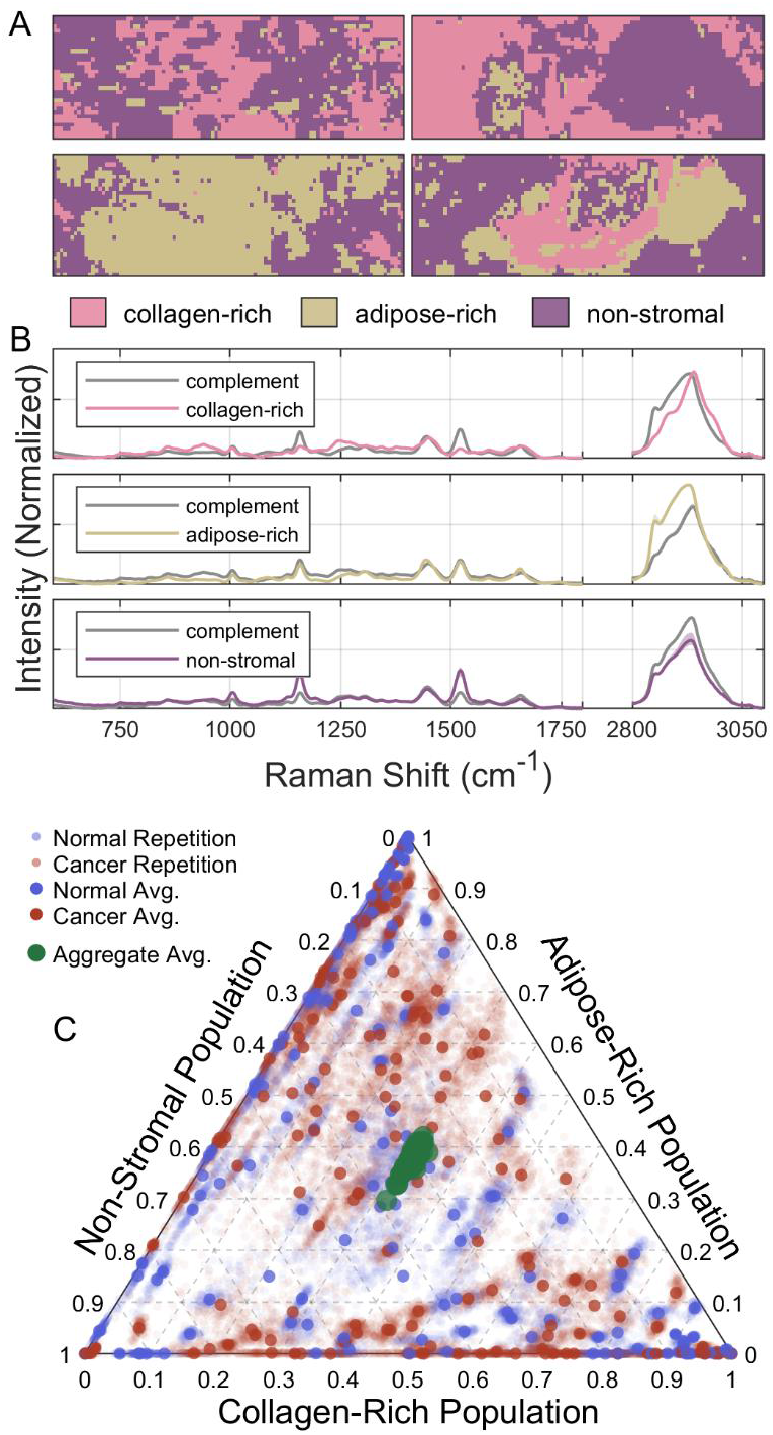
Posterior evaluation of label propagation for HBT modality segmentation. A) Spatial mappings of the most-commonly assigned collagen-rich (pink), adipose-rich (brown), or non-stromal (purple) spectral segments across 96 LP repetitions at each pixel in the 4 Raman images illustrated in Fig. 1. B) Averaged (solid lines) spectra and their IQRs (shading) of each segment and its complement across the 96 LP reps. Complement spectra are shown in gray color while collagen-rich (pink), adipose-rich (brown), and non-stromal (purple) segments are shown from top to bottom. C) Segment populations within RHIs are depicted in a ternary plot. Fractions of the 3000 spectra within an image assigned to each spectral segment within a particular LP repetition are represented as small red (cancer) or blue (noncancer) markers. Larger red (cancer) or blue (noncancer) markers represent average segment populations for each individual RHI across all 96 LP reps. Large green markers represent the average segment populations aggregated across all RHIs for each of the 96 LP reps.

### Artificial Neural Networks (ANNs) for Cancer/Noncancer (C/NC) Classification of Raman Hyperspectra

Fig. 3 illustrates the construction of the cancer/noncancer models. C/NC models are artificial neural networks (ANNs) that are combinations of convolutional neural networks (CNNs)^53^ for feature extraction and deep learning networks (DLNs)^54^ for binary C/NC classification. The ANNs take Raman hyperspectra as inputs and output posterior class weights of both cancer and noncancer for each spectrum. In addition to deeper CNNs like residual networks^55^, the architecture used in the CNN was inspired by the discrete wavelet transform (DWT), a foundational method for local frequency-domain decomposition of a discrete input signal^56,57^. Like the DWT, the CNN implementation is composed of distinct convolutional blocks that recursively convolve an input signal with a digital filter and then downsample to reduce resolution. After the CNN, concatenated outputs are input to a DLN to return posterior weights for subsequent use in spectral classification. Detailed architectures and hyperparameters of CNN blocks and the DLN are included in the Appendix. For training and testing, the set of 957,000 spectra in 319 RHIs is split by image with a train-to-test ratio of approximately 80% to 20%. Spectral preprocessing is described in the Appendix. First, 32 cancer and 32 noncancer testing images are selected at random as a class-balanced test set, and the remaining 255 images comprise the training set. Undersampling of the larger training class (cancer histology) is necessary to ensure class balance. In total, 468,000 spectra (372,000 training and 192,000 testing) from each class are used for testing and training. Supporting Fig. S2 shows the accuracies and cross-entropy losses of representative training and validation sets versus training epoch.

**Figure 3.**
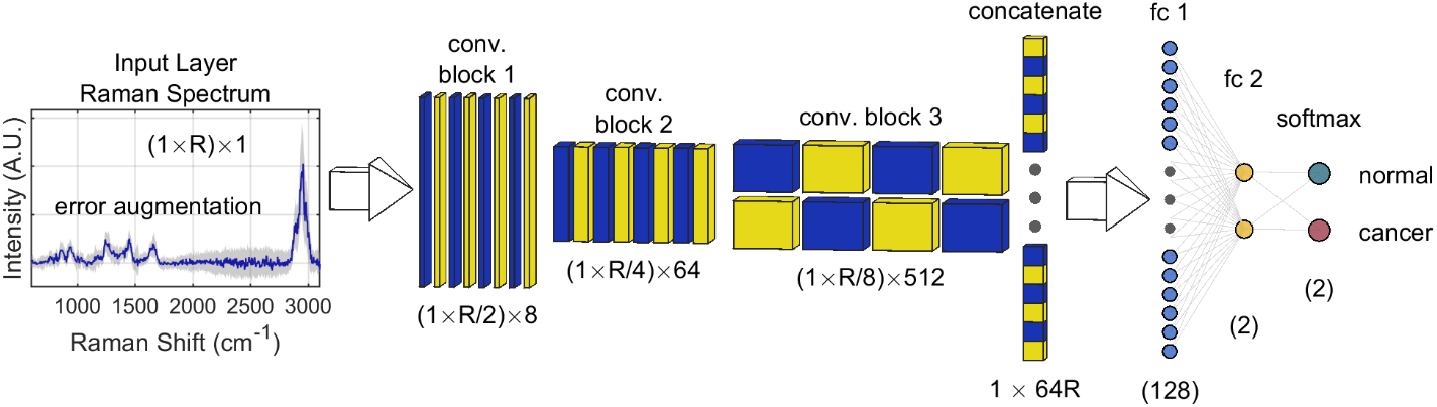
Cancer/Noncancer classification model for Raman hyperspectra. From left to right, inputs are 1-dimensional Raman spectra containing *R* Raman shift increments. Input training spectra first undergo augmentation according to estimated photon-counting error, as described in the Appendix. Augmented spectra are passed through a CNN containing 3 convolutional blocks. Each block contains 8-kernel convolutional layers followed by batch normalization and ReLu activation layers. Finally, a maximum pooling layer is used to reduce dimensionality. Convolutions are implemented with strides of 1 and kernel sizes of 2, while max. pooling is implemented with strides and pool sizes of 2. For simplicity in notation, the number of Raman shift increments *R* as displayed must be a factor of 2, and convolutional layers are zero-padded such that each CNN block results in exactly 1/2 the number of input increments. After the CNN, all output increments are passed through a concatenation layer and into a DLN for classification. The DLN outputs 2 scores, corresponding to posterior weights of the cancer and noncancer classes for the input spectrum. The output layer is preceded by a 2-layer fully connected network, with the first layer reducing dimensionality to 128 increments and a second reducing to two (2). Batch normalization and a ReLu activation layers follow fc 1, and a softmax layer converts the outputs of fc 2 to posterior class weights. ANN architecture is completely detailed in the Appendix.

Because many unique test/train splits are possible in a data set of this size, a single split cannot be expected to sufficiently describe the range of outcomes that may occur in a more general situation. Many methods exist to estimate such ranges of outcomes, such as *k*-fold cross validation or bootstrap aggregation. Here we use repeated random subsampling (RRSS), also known as Monte Carlo cross-validation^58^, in which randomized repetitions (reps) of the test/train split procedure are implemented, generating multiple realizations of training and testing data. C/NC models are trained independently of one another using each training set rep, then are tested using the corresponding testing set rep. Training spectra are error-augmented at the start of each training epoch by random sampling of their intensities from normal error approximations, whose magnitudes were obtained numerically through repeated denoising as described in the Appendix. Such augmentation allows intensities to fluctuate according to their measurement errors, which avoids overfitting spectral class differences that are smaller in magnitude than Poisson-distributed photon-counting errors. Testing results in a pair of posterior class weights for each of the 192,000 spectra. In total, 512 RRSS reps were produced, which are analyzed *a posteriori* to obtain classification qualities (CQs), predicted class spectra, and spatial mappings of cancer index (CI), which are isolated posterior weights of the cancer class for each spectrum.

## RESULTS AND DISCUSSION

### Posterior Evaluation of C/NC Classification of Raman Hyperspectra

Fig. 4 contains posterior evaluation of C/NC model implementation with RRSS. Figs. 4A and 4B evaluate classification qualities (CQs) of each of the 512 repetitions (reps) obtained using RRSS, while Figs. 4C and 4D evaluate spatial CI properties and predicted spectral properties in comparison to ground truth (GT) histopathological assignments, respectively. In Fig. 4A, each rep is represented with a single marker indicating accuracy and corresponding area under its receiver operating characteristic (ROC) curve (AUROC). AUROCs represent the areas under the ROC curves that are displayed in Fig. 4B for all 512 reps, and accuracies represent the ratio of the number of correct classifications to the total number of spectra in the test set of each rep (192,000). While the AUROC is independent of threshold, accuracies require the use of thresholds on the CIs such that spectra with CIs greater than the threshold are positive cancer predictions and those less than or equal to the threshold are negative predictions. Thresholds are chosen for each RRSS rep such that accuracy is maximized. Accuracies and AUROCs show no observable dependence versus corresponding thresholds (supporting Fig. S3). Figs. 4A and 4B show CQs to be consistent across reps, returning median accuracy of 0.846 and AUROC of 0.913, suggesting reasonable distinction of cancer and noncancer spectra. Particularly good agreement is observed among RRSS models, as all reps return accuracies and AUROCs that are within approximately 10% of the median values. ROC curves (Fig. 4B) yield similar observations as all curves display consistent characteristics, suggesting robustness of the C/NC models against variations in the training and test sets and reproducibility of CQ.

**Figure 4.**
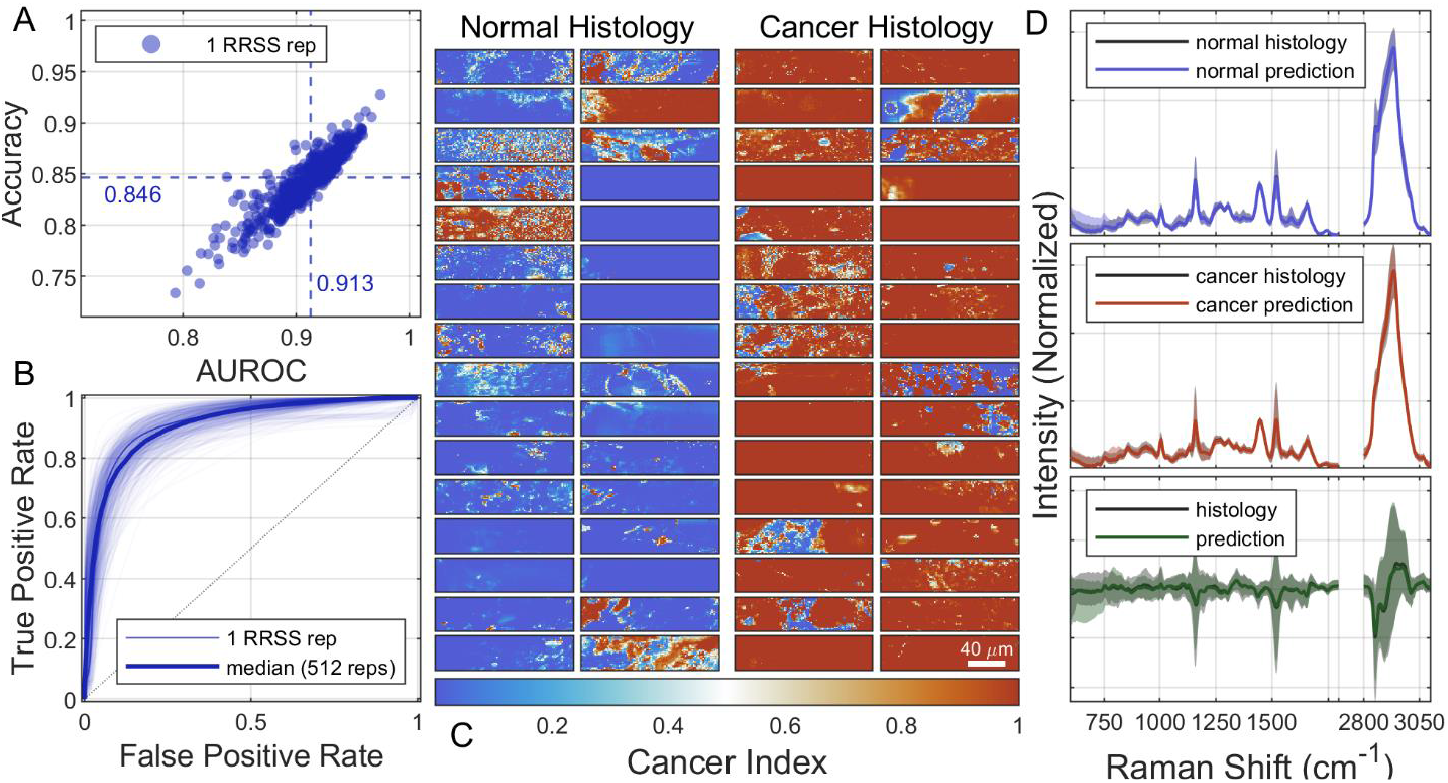
Posterior Evaluation of C/NC models. A) Each marker represents two classification quality (CQ) descriptors, accuracies and areas under the receiver-operating-characteristic curves (AUROCs), of a single implementation of RRSS and a C/NC model. The median accuracy (0.846) and AUROC (0.919) across all 512 spectral C/NC classification reps are indicated as horizontal and vertical blue, dashed lines, respectively. B) ROC curves for RRSS reps, plotting true vs. false positive rates from which the AUROCs in A) were computed, are shown as thin blue lines, while the median behavior over all reps is indicated in a thicker blue line. C) Heatmaps showing spatial distributions of cancer index (CI) in a representative RRSS rep (accuracy 0.878, AUROC 0.932), containing 64 RHIs (32 C, 32 NC). RHIs from samples with normal histology are in 2 columns to the left and with cancer histology to the right. CI is indicated in the color scale below the heatmaps, with blue colors representing low CI and red representing high CI. D) Predicted noncancer (blue) and cancer (red) average spectra are compared to their ground truth (GT) histological predictions (gray) in the upper and center panels, while the cancer-noncancer (C-NC) difference spectra, in which the noncancer average spectra are subtracted from the cancer average, predicted by the C/CN models (green) are compared to their GT (gray) in the lower panel. In all panels, solid lines represent average spectra while shading represents IQRs over 512 RRSS reps.

Fig. 4C shows spatial CI maps returned by the ANNs for a representative RRSS rep near the median accuracy and AUROC. Normal tissues are in 2 columns on the left and cancer tissues in 2 columns to the right. The diverging color scale below indicates low CI and high CI in blue and red colors, respectively. Although many images on the left are mostly blue and many on the right mostly red indicating high CQ in these images, some images display disagreement between CI and histopathology that is persistent across their entire areas, most likely indicating labeling errors or latent features in histopathological assessments. In other images, spatially contiguous groupings of similar CIs exist, but CI varies in groups across the images, suggesting that cancer recognition by the C/NC model may be dependent on tissue modality. Predicted average spectra for the cancer (red, upper panel) and noncancer (blue, center panel) are compared to their GT (C/NC histopathology) spectra (black) in Fig. 4D above their corresponding difference spectra, in which the predicted (GT) noncancer spectrum is subtracted from the predicted (GT) cancer spectrum for each RRSS rep. In all panels of Fig. 4D, solid lines indicate average spectra while corresponding IQRs over 512 reps are shown as shaded regions in corresponding colors. Upper and center panels of Fig. 4D show clearly that predicted spectra provide good recovery of GT cancer and noncancer spectra, leading to good recovery of the cancer-noncancer (C-NC) difference spectra as well. Interestingly, C-NC difference spectra indicate negative intensity at Raman shifts corresponding to symmetric and asymmetric CH_2_ stretching (approximately 2853 and 2896 cm^-1^) and at Raman shifts corresponding to carotenoids (approximately 1519, 1156, and 1004 cm^-1^), which suggests the models recognize decreases in both lipid and carotenoid intensities in cancerous tissues, in agreement with the phenotypical properties of CAAs and FC that display decreases in lipid prominence in cancerous tissue^30^. Decreases in carotenoid intensity may indicate a decrease in carotenoid prominence, or that the cancerous tissue is in a state of oxidative stress, leading to oxidation of carotenoids by excess ROSs.

### C/NC Classification in Isolated Spectral Segments

It is informative to evaluate C/NC model performance within the collagen-rich, adipose-rich, and non-stromal spectral segments approximating HBT modalities obtained using label propagation (LP). Spectral segments are isolated from one another either before or after C/NC model training, and C/NC behaviors such as CQs and C-NC difference spectra are evaluated within spectral segments. Two orders of operation are possible: 1) perform C/NC classification on non-segment-isolated spectra first, then partition resulting CIs according to their LP segmentation labels and compute CQ, or 2) isolate spectral segments before C/NC classification and train segment-isolated C/NC networks using only spectra that were assigned to a particular spectral segment to obtain CIs and subsequently CQ. RRSS and train/test splitting in the latter case are performed as described above with adjustment for differing numbers of spectra, obtaining segment-isolated CIs for 512 RRSS reps. For clarity, the former and latter sequences are given distinctions of non-isolated and segment-isolated C/NC classifiers, respectively.

Fig. 5A shows CQs obtained from segment-isolated C/NC classifiers as filled markers, while CQs obtained from non-isolated C/NC networks are shown as shaded circles. The panels of Fig. 5A show clearly that there is little difference in CQ between the segment-isolated and non-isolated C/NC networks after C/NC classification. Median accuracies and AUROCs show no significant differences and significant overlap among the dispersion of CQs between the two approaches is observed. This suggests that, despite the training set containing spectra originating from multiple HBT modalities, non-isolated C/NC networks learn multimodal class distinctions comparable to segment-isolated networks in which the training and testing sets originate from a single spectral segment and a narrower range of HBT modalities. Furthermore, tight CQ grouping is again observed among the 512 reps, indicating robustness against training and test set variability and reproducible CQ. Overall, weaker CQ is observed in the collagen-rich and non-stromal segments. A possible source for weaker CQ may lie in biological variability, as both collagen-rich and non-stromal spectral groups may contain contributions from multiple HBT modalities. Another possible source is lower signal-to-error ratio (S/E), as lipids in adipose modalities have large scattering cross-sections among important biomolecules and therefore yield spectra with relatively high Raman intensities^34^. The highest observed CQs originate from the adipose-rich segment, with a median AUROC of 0.953 and accuracies often exceeding 90%, indicating robust and consistent C/NC models. These results demonstrate reliable distinction of cancerous adipose tissue from noncancerous, indicating consistent distinction of Raman spectra of CAAs from those of normal adipocytes. This strongly suggests that detection of the cancer field effect with Raman microscopy is possible, as reliable CQ is obtained in detection of cancerous stromal tissue in regions that are nearby excised tumors, particularly in adipose-rich regions.

**Figure 5.**
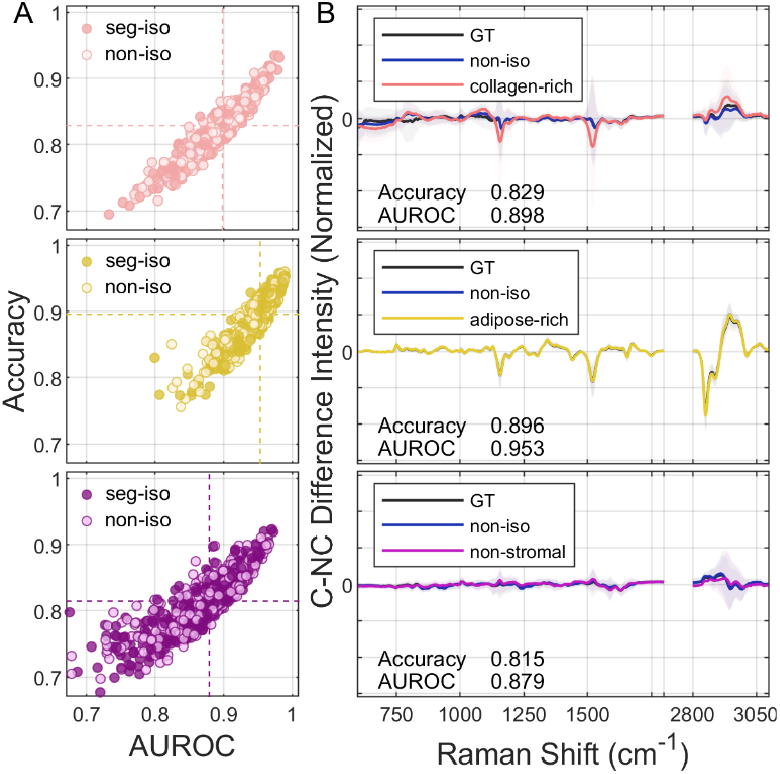
Posterior evaluation of C/NC classification within spectral segments. A) Each marker represents the accuracy and AUROC obtained from a single RRSS rep. Reps obtained using segment-isolated C/NC classifiers are filled markers while reps using non-isolated classifiers are shaded circles. The median accuracies and AUROCs are indicated as horizontal and vertical dashed lines in corresponding color. Reps for the collagen-rich segment are shown in the upper panel, adipose-rich in the center panel, and non-stromal in the lower. B) C-NC difference spectra for GT (black), non-isolated (blue), and segment-isolated classifiers (pink, yellow, purple). Collagen-rich difference spectra are displayed in the upper panel, adipose-rich in the center panel, and non-stromal in the lower panel. Solid lines indicate mean difference spectra while shading represents IQRs over all 512 reps.

C-NC difference spectra for each segment-isolated classification are shown in Fig. 5B, with collagen-rich spectra in the upper panel, adipose-rich spectra in the center panel, and non-stromal spectra in the lower panel. In each panel the GT C-NC spectrum is shown in black color, while those of non-isolated C/NC classification are shown in blue color. Each of the C-NC spectra of segment-isolated C/NC networks are shown in pink, yellow, or purple color for collagen-rich, adipose-rich, and non-stromal spectral segments, respectively. Solid lines indicate median difference spectra and shaded regions represent IQRs over all 512 reps. C-NC spectra in Fig. 5B indicate good recovery of GT cancer and noncancer spectra and spectral differences, as both non-isolated (blue) and segment-isolated (various colors) predictions are in good agreement with the GT C-NC difference spectra (black). The most consistent spectral recoveries are obtained in the adipose-rich segment, reflecting the higher CQ that was observed in this segment. Adipose-rich C-NC spectra show reduced lipid intensity in adipose-rich cancer tissues, again (*c*.*f*. Fig. 4) consistent with reliable recognition of CAAs and normal adipocytes. Reduced carotenoid intensity suggests either a reduction in carotenoid prominence or an increase in oxidative stress as cancer progresses. Slight reduction of carotenoid is also observed in cancerous collagen-rich tissue, especially by the segment-isolated C/NC classifier, further suggesting recognition of field cancerization through oxidative stress, reflecting the conversion of normal fibroblasts to CAFs through increased prominence of ROSs in the fibrous ECM modality.

### Improved CQ Through Exclusion of Hyperspectra with Low Signal-to-Error (S/E) Ratios

Improvement of CQ through exclusion of testing spectra having S/E lower than specified thresholds is explored using the non-isolated C/NC networks, which have returned consistent and robust multimodal class recovery. That is, noisy, low-quality test measurements are discarded and only those measurements that meet or exceed a certain quality are evaluated. S/E is the average of the ratio of spectral intensities to their corresponding errors that arise from photon-counting taken over all Raman shifts in the spectrum. We note that the S/E range across the data set is broad, with a minimum of just 0.35 versus a maximum S/E of approximately 125. S/E thresholds were chosen as quantiles of the S/E estimates over all 957,000 spectra, such that there are 20 thresholds in 5% increments between the 0^th^ and 95^th^ percentiles, resulting in 20 spectral groups that contain equivalent numbers of spectra. The fraction of spectra that are excluded therefore begins at zero for the first S/E threshold (S/E≥0) and increases in 5% increments until the last threshold, in which 95% of spectra are excluded. Fig. 6A displays occurrence fractions across the reps of each spectral segment in test spectra that meet or exceed indicated S/E thresholds. Markers and solid lines indicate average occurrence fractions, while shaded regions indicate IQRs over all reps. For example, all test spectra are included at the first S/E threshold, and occurrence fractions represent average populations of the collagen-rich (pink), adipose-rich (yellow) and non-stromal (purple) spectral segments across all spectra.

**Figure 6.**
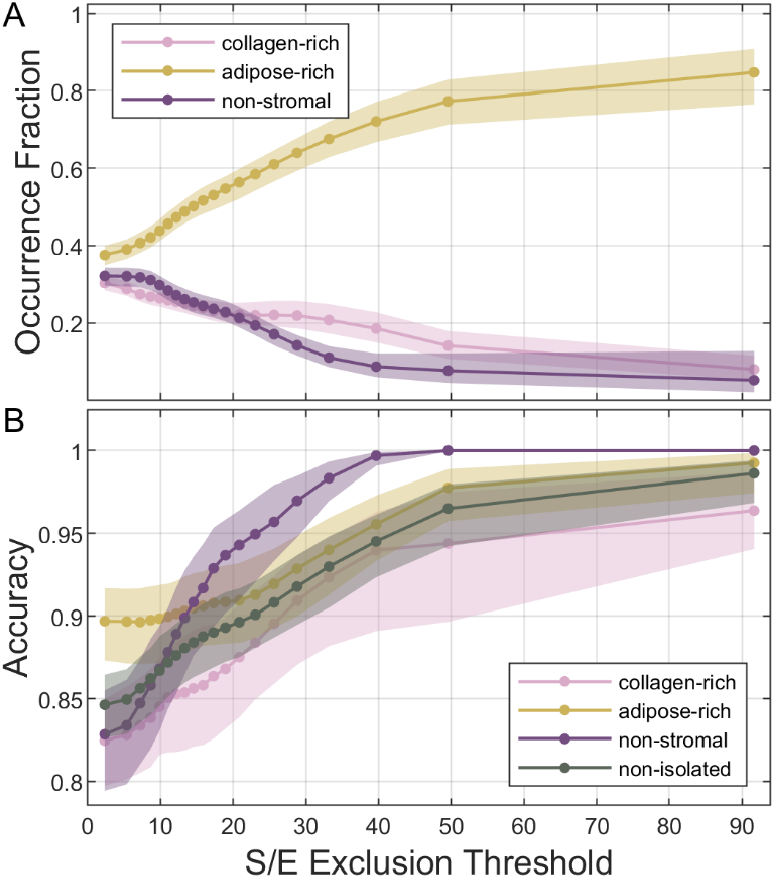
C/NC classification vs. signal-to-error ratio (S/E). A) Occurrence fractions of each spectral segment in sets of test spectra that have average S/E greater than or equal to a specified threshold. B) Accuracy is shown to increase as spectra with lower S/E are excluded from testing. In both panels, solid lines and markers represent averages and shading represents IQRs over 512 RRSS reps. Collagen-rich occurrence fractions and accuracies are shown in pink color, adipose-rich in yellow, and non-stromal in purple. Non-isolated accuracies are shown in green color in panel B).

As the S/E threshold is increased and lower S/E spectra are excluded, larger fractions of the remaining spectra originate from the adipose-rich segment, indicating that higher S/E spectra are obtained from adipose modalities in HBT as opposed to non-adipose modalities. Fig. 6B shows accuracies versus S/E exclusion threshold, both isolated within each spectral segment and non-isolated spectra, with collagen-rich stromal accuracies in pink color, adipose-rich accuracies in yellow, and non-stromal in purple. Non-isolated accuracies vs. S/E exclusion threshold are shown in green color. Markers and solid lines represent median accuracies while shaded regions indicate IQRs over all 512 reps. Fig. 6B shows CQ to increase dramatically with increasing S/E threshold in all spectral segments, approaching pure distinctions at the highest S/E exclusion thresholds. This predictably suggests that C/NC models perform best on high quality spectra with high S/E and conversely return inferior CQ when encountering spectra of lesser quality at lower signal-to-error ratios. While the CQ increase is observed versus increasing S/E in all spectral segments, as indicated in Fig. 6A, collagen-rich and non-stromal segments are poorly represented at higher S/E, suggesting that high CQs largely originate from reliable discrimination of Raman hyperspectra of normal adipocytes from those of CAAs, an observation of field cancerization in human breast tissue.

### Larger Physical Acquisition Areas Increase S/E but Not CQ

Because higher CQ is obtained using higher quality spectra with higher S/Es, we explore increasing S/E through block-averaging of the Raman hyperspectra over spatially contiguous regions spanning larger areas of the sample, such that S/E is increased through collection of larger numbers of photons. Spatial blocks are chosen to be square areas containing integer numbers of acquisition pixels for simplicity. Including the acquisition sizes, we designate 9 square block sizes ranging from 1.5 to 30 *μ*m in length along one side. For block-averaged spectra, non-isolated C/NC networks and test/train RRSS are performed in the same manner as described for spectra at the acquisition size, adjusting for changes in the numbers of spectra accordingly. Additional translational and rotational augmentations are used to generate training spectra for the block-averaged C/NC networks, as detailed in the Appendix. As the notion of spectral segment is obscured through possible averaging over multiple LP labels within single spectral blocks, only non-isolated C/NC networks are evaluated. S/E thresholds and exclusions are performed in the same manner as described in discussion of Fig. 6.

Fig. 7 shows accuracies versus S/E exclusion thresholds. The expected increase in S/E versus block size is apparent in Fig. 7A, as larger block sizes result in larger S/E. However, it is also clear that increased block size does not result in higher CQ, as accuracies obtained at the highest S/Es decrease versus increasing block size. Although contradictory to the expected increase in CQ with increased S/E, this result implies that biological fluctuation significantly contributes to the resolution of cancer/noncancer tissue in Raman hyperspectra. Because spatial acquisition areas are larger, mixing of spectra arising from multiple HBT modalities inevitably occurs, which appears to blur classification boundaries between cancerous and noncancerous spectra, leading to poorer CQ at larger block sizes and higher S/E. That is, these results clearly demonstrate that assessment of C/NC in Raman hyperspectra on smaller physical scales with lower S/E and higher purity in HBT modality leads to better C/NC distinction by the ANNs than C/NC assessment on larger physical scales using averaged measurements in which multiple tissue modalities are mixed. Rather, CQ appears to follow a consistent pattern versus increasing S/E quantiles, as indicated more distinctly in Fig. 7B, showing poorer CQ at lower S/E quantiles and increasing as S/E quantile increases. This clearly suggests that the contributions of HBT modality mixing within spectra spanning larger spatial areas, i.e., biological fluctuations, are significant in C/NC distinction using Raman hyperspectra, as larger block sizes worsen C/NC model performance.

**Figure 7.**
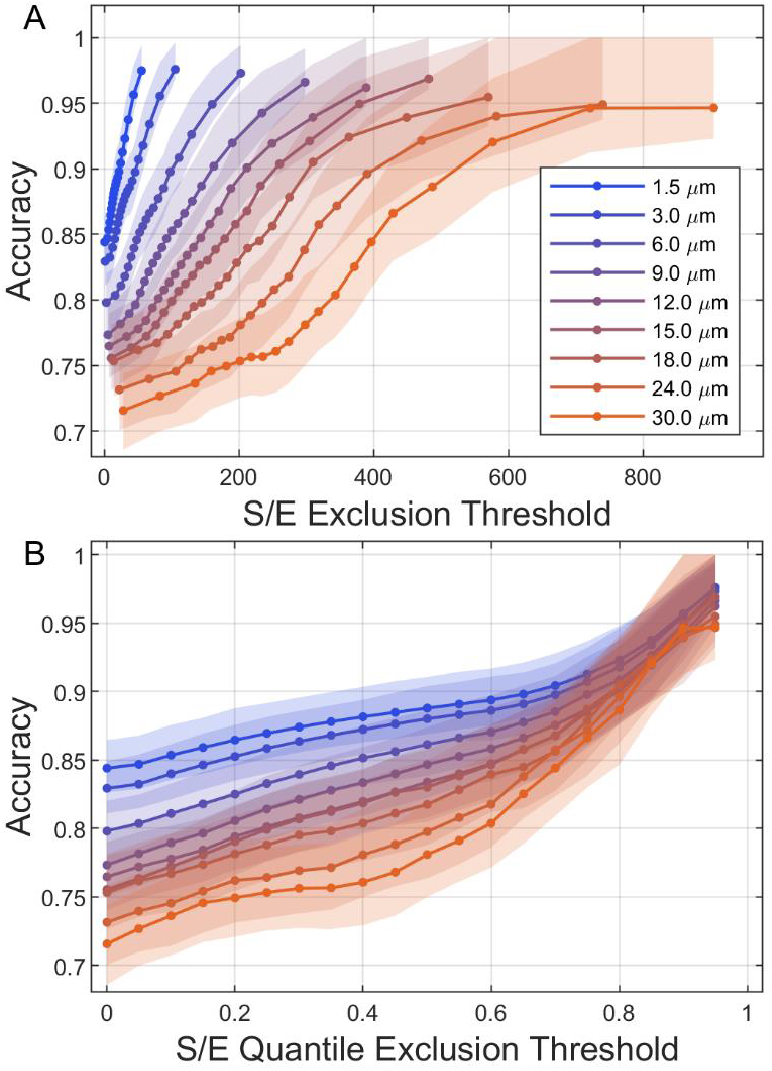
Accuracies of C/NC classification models versus S/E exclusion threshold and physical spectral acquisition size. A) Accuracies versus S/E exclusion threshold for increasing block sizes. Smaller block sizes appear in blue colors while larger sizes appear in orange. S/E thresholds on the horizontal axis are 5^th^ percentiles (0-95%) of the S/E observations at that block size. B) The same accuracies and block sizes shown in panel A), adjusted such that S/E quantiles (0-95%) are represented on the horizontal axis.

## OUTLOOK

Reproducible detection of field cancerization in multiple modalities of human breast tissue, in particular, detection of converted cancer-associated adipocytes, using Raman hyperspectral imaging and machine learning, artificial neural network models has been demonstrated. This articulates the potential of medical devices using Raman spectral data for fast and accurate cancer/noncancer diagnosis in cancer patients. However, highly accurate detection is only possible at very high signal-to-error ratios due to the contributions of measurement errors in lower quality data. Although S/E can be improved through acquisition of the Raman spectra such that larger regions of the sample are represented, mixing of differing HBT modalities over larger regions gives rise to significant biological variation, negating the benefits of increasing the measurement quality. In conclusion, although Raman hyperspectral images show great potential in providing detailed biochemical representations of complex biological systems, data quality remains a substantial issue that must be improved before practical use is possible. In terms of the detection of field cancerization, RHI offers a highly attractive alternative to intraoperative histopathological diagnosis of margin tissues, and, with further development, also offers the potential for surgeons to test tumor margins *in situ* without removing or damaging more tissue than necessary, potentially benefitting tissue-conserving surgeries in the treatment of many cancers.

## ACKNOWLEDGEMENTS

We would like to thank Noriko Ishida for laboratory preparations. This work was partially supported by AIST-Osaka University Advanced Photonics and Biosensing Open Innovation Laboratory (PhotoBio-OIL), and Japan Science and Technology Agency (JST) COI-NEXT Grant Number JPMJPF2009.

The authors declare no competing interests.

## APPENDIX: Detailed Methods

### Preprocessing

Spectra are preprocessed to remove electronic bias according to manufacturer specifications, and for cosmic rays with outlier detection (+8σ deviation from mean intensity), before denoising with singular value decomposition (SVD)^59^. SVD is used to decompose each image into its singular values (SVs). Insignificant SVs having magnitudes less than 1% of the maximum SV magnitude are set to zero, and the image is reconstructed using the remaining 12 SVs. Poisson-distributed photon-counting errors are estimated to be normal and are propagated numerically with repeated random sampling of each intensity and subsequent SVD denoising with keeping the 12 SVs. Each intensity measurement in an image first undergoes random sampling according to a normal distribution having mean and variance equal to the observed intensity. Next, the image is denoised using SVD and stored. Denoised spectra are then estimated as mean intensities over 256 repetitions of sampling-denoising, and the errors as the standard deviation over the same intensities. Signal-to-error ratios (S/Es) are then estimated for each spectrum through pointwise division of the spectral intensities at each Raman and propagated errors. The mean S/E representing the spectrum is the average of the ratio over all Raman shifts between 600 and 3100 cm-1. Each spectrum is transformed to a common Raman shift axis using cubic Hermite polynomial interpolation. Next, an extrinsic background spectrum (EBS)^45^ is estimated through identification of the least intense set of spectra across all Raman images. The EBS is then used as the linear component in a baseline correction using asymmetric least-squares polynomial regression^60^, allowing for linear combination of the EBS with other components of baseline that may arise from, e.g., autofluorescence in the HBT. Each baseline-corrected spectrum and associated errors are normalized by sum intensity of in the 600-1800 and 2800-3100 cm^-1^ ranges.

### Label Propagation

Label propagation (LP) makes use of a partial set of labeled data to propagate labels throughout unlabeled members of the data set. We use Raman intensities of prominent molecules to generate sets of initial labels. For example, adipose tissue shows high CH_2_ stretching intensities near 2850 cm^-1^ and fibrous collagen tissues of the ECM show high amide III stretching intensity near 1250 cm^-1^. Initial labeling is performed in the intensity space for adipose and collagen target molecules using sum intensities in the band of Raman shifts of interest, e.g., 2840-2860 cm^-1^ for adipose and 1240-1260 cm^-1^ for fibrous collagen, such that spectra having intensities less than the 1st percentile intensity across all spectra in all images are labeled as non-target spectra and those with intensities greater than the 99th percentile are labeled as adipose or collagen target spectra. The remaining 98% of spectra remain unlabeled. Spectra are transformed to a principal component (PC) space such that full spectral information is utilized, where each spectrum carries its initial label, and the PCs of the spectra having non-adipose and adipose labels are used to train a k-NN classifier using significant PCs having more than 95% explained variance. Unlabeled spectra are assigned posterior weights for the target or non-target classes, which are ratios of the number of neighbors in each class to the total number of neighbors. Posteriors are computed in localized neighborhoods for each unlabeled spectrum from the labels of the nearest *k* neighbors in the training set, with *k* being the square root of the number of spectra in the training set. New labels are generated for unlabeled spectra such that those having 100% of their neighbors in the target set are labeled as target spectra, those having 0% of their neighbors in the non-target set are labeled non-target spectra, while spectra having mixed posterior weights remain unlabeled. Spectra that gain new class labels are added to the training set for the next iteration, in which a new k-NN model is trained on the updated training set. Random undersampling of the larger class in the training set is used to maintain class balance in subsequent iterations. New labels are assigned to unlabeled spectra, and the training set is once again updated with new members. This process continues until the set of labels has stabilized. Practically, this occurs when the number of spectra that remain unlabeled converges, changing less than a specified threshold (0.01%) over iterations. Spectra with adipose or collagen labels that are members of the training set in the convergence iteration are assigned adipose or collagen labels, while non-adipose-labeled and remaining unlabeled spectra are assigned non-adipose labels.

### Artificial Neural Networks (ANNs)

In each CNN block, a convolutional layer is followed by a batch normalization, a ReLU activation layer, and a maximum pooling downsampling layer. Convolutional layers are implemented with window sizes and strides of 2 increments and 8 convolutional kernels. Maximum pooling layers that follow are also implemented with strides of 2 increments, downsampling the input by a factor of 2 in each convolutional block. Following the CNN blocks, outputs are passed through a concatenation layer and into a deep-learning network (DLN) for classification. Concatenated outputs are first subject to a fully connected layer, reducing the representation to a size of 128 increments. Output increments are then passed through a batch normalization then a leaky ReLU activation layer. A second fully connected layer reduces the representation to two scores representing each class, which are then converted to posterior class weights using a softmax layer.

All 957,000 spectra in 319 images are split by image into test and training sets with a train-to-test ratio of approximately 80% to 20%. First, ensuring that class sampling is balanced in the test set, 32 cancer and 32 noncancer testing images are selected at random. The remaining 255 images comprise the training set. Because class sampling is slightly imbalanced across the data set (51.1% cancer samples vs. 48.9% normal tissue samples), undersampling is performed on the training set at the spectral level. Of the 255 images in the training set, 131 images are cancer samples and 124 are normal, resulting in 393,000 cancer spectra and 372,000 normal spectra. 21,000 cancer spectra are discarded randomly without regard to image of origin, balancing sampling in the training set at 372,000 spectra per class. 468,000 cancer and 468,000 normal spectra are included in the final test/train split, with a total of 192,000 (91,000 cancer and 91,000 normal) spectra in the test set and 744,000 (372,000 cancer and 372,000 normal) spectra comprising the training set.

A C/NC model is trained using the resulting training set. Training is initiated with a mini batch size of 128 spectra and a learning rate of 1×10^-3^. At the start of each epoch, training spectra are augmented using spectral errors as estimated from the SVD denoising process described in Preprocessing. This allows the model to avoid overfitting spectral differences among classes that are smaller than the error magnitudes allow. Training continues for (24) epochs, after which the models were sufficiently converged in their optimization of the cross-entropy loss. A demonstration showing convergence of classification qualities and cross-entropy losses versus epoch in the training set and a validation set for a representative model is shown in supporting Fig. S2. Testing is performed using the trained C/NC model. Each of the 192,000 test spectra are passed through the trained model, yielding 192,000 cancer/noncancer posterior weight pairs. Because the posterior pairs are normalized, we isolate the cancer posterior, termed cancer index (CI) for simplicity, for assessment of classification quality and other relevant quantities and properties, such as predicted class spectra.

Adjustable parameters of the CNN and DLN were optimized using adaptive moment estimation^61^ such that the objective function, the cross-entropy loss between the predicted and true labels in the training set is minimized. Hyperparameters were chosen such that minimal values that achieved stable CQs were selected as optimal. Minimal kernel lengths of 2 increments and dyadic strides were used in both convolution and pooling, resulting in a 2x downsampling in the number of increments for each convolutional block. The number of blocks was chosen using the elbow method, i.e., subsampled test/train splits were tested initially with a single block, then with 2 blocks and so on, integer-wise until classification quality stabilized. CQ stability was achieved with 3 convolutional blocks, with insignificant CQ improvement being observed with 4 blocks. The number of kernels in each of the 3 blocks was increased integer-wise until CQ achieved stability. Similarly, the DLN was initialized with batch normalization and ReLU layers followed by a single, binary fully connected layer and a softmax output layer. An additional fully connected layer was added to the front on the DLN, and the size of the additional layer was increased until CQ stability was achieved.

### Augmentation

During training, normalized spectra are error augmented by random sampling each intensity from normal distributions with means and standard deviations being the intensities and associated errors estimated through numerical error propagation (see Preprocessing). Augmentation is performed on every spectrum in the training set at the initialization of each training epoch. In addition to error augmentation, translational and rotational augmentations were performed during training for block-averaged spectra. Each RHI is 30×100 grid of Raman spectra, and block-averaged spectra are produced by averaging the spectra within subsections of the grid. Although a certain number of distinct, non-overlapping blocks of a given size are possible within the grid, translations smaller than the block size produce unique observations. Furthermore, because the orientation of the measurement is arbitrary, rotations also result in unique observations. At the initialization of each epoch, the number of distinct blocks possible in the 30×100 grid is subsampled randomly from each image from the set of all possible translated blocks, allowing spatial overlap among augmented blocks. Each block is then rotated randomly in a range of 0-360 degrees. We note that raw block sizes must be a factor of √2 larger than the target block size, allowing for capture of the diagonal axis of the blocks within the selected block size. Block-averaging is then performed on each block producing block-averaged spectra that have been augmented to represent measurement error, as well as translation and rotation of the measurement orientation.

### Repeated Random Subsampling

RRSS, also known as Monte Carlo cross-validation^58^ is an approach that repeatedly implements the test/train split procedure described above (see ANNs) to generate multiple test/train splits which are then treated as independent models. In total, 512 test/train splits, or RRSS repetitions (reps), were produced for this work, resulting in 512 unique models by which to compare model outputs. For example, 512 reps produce 512 classification accuracies, 512 ROC curves, 512 sets of predicted class spectra, etc., which can then be analyzed *a posteriori* to interpret model quality and effectiveness.

### Computational Hardware

All computations were performed on a desktop computer equipped with 128 GB RAM, a 3.2 GHz Intel Core-i9 CPU, and Nvidia 3090 GPU with 24 GB onboard RAM. Custom software used MATLAB R2024a and associated toolboxes.

### Experimental Procedures

Normal and malignant tissue samples were removed from patients undergoing partial mastectomies for treatment of breast cancer after obtaining informed consent. Patient identities and other identifying information was kept private from the researchers. Tissues are stored at −80°C prior to experimental preparation. Hematoxylin and eosin staining was performed on a portion of each sample while other portions are preserved for Raman measurement. Histological images were obtained for each sample and were assessed for malignancy by a qualified histopathologist. Samples were thinly sliced with approximately 10 *μ*m thickness, placed on a quartz substrate and immersed in PBS buffer in preparation for Raman measurements. Optical measurements were performed using a self-made, line-illumination Raman microscope^34^. Excitation light with wavelength 532 nm and irradiance was 5.5 mW/μm2 was focused through a water immersion objective lens with 60× magnification and 1.27 numerical aperture (CFI SR Plan Apo IR 60×C WI, Nikon). Photons scattered from the sample were collected through the same objective lens and passed through a spectrophotometer (MK300, Bunkoukeiki), where they were detected by a cooled CCD camera (PIXIS 400B, Teledyne Princeton Instruments). Measurements were performed such that environmental conditions were consistent for each sample, using procedures established for biological samples^34^.

## SUPPORTING FIGURES

**Figure S1.**
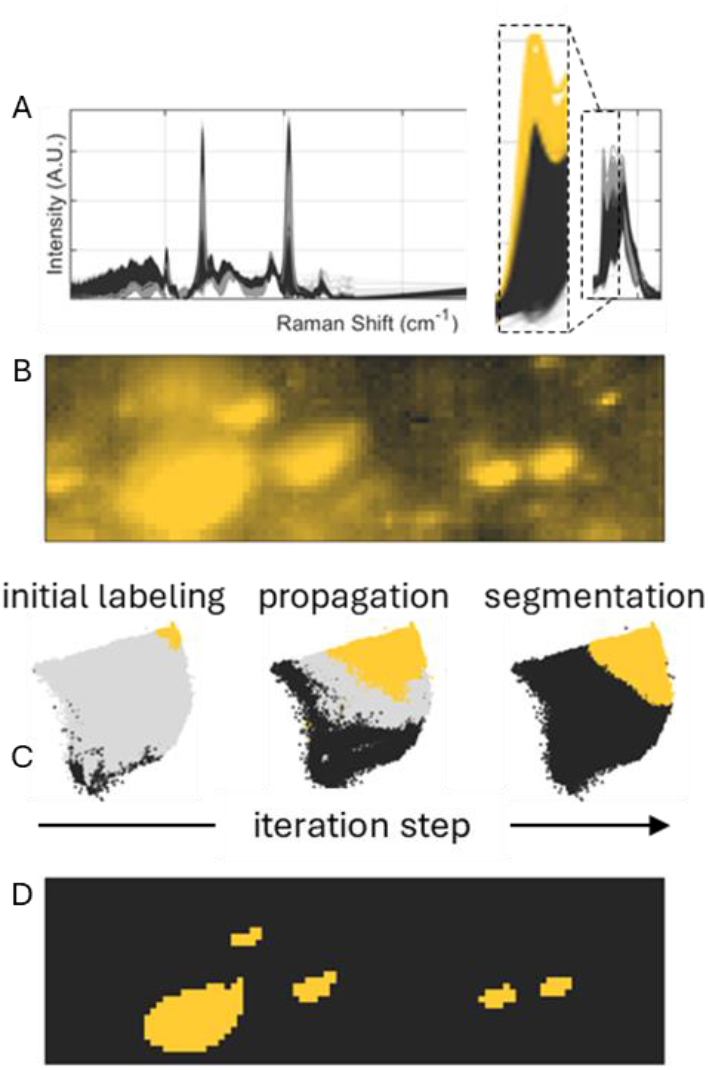
Label propagation for segmentation of adipose-rich tissue modalities. A) Spectral properties used for initial labeling using the sum intensity in the band of Raman shifts between 2840 and 2860 cm-1. B) Spatial mapping of intensities depicted in A). C) (left) Spectra are transformed to an initial PC space with initial labels, in which adipose-rich spectra are shown as yellow markers, non-adipose spectra in black, and gray markers indicate spectra that remain unlabeled. (center) Intermediate labels during propagation. (right) Convergence of LP-PC space. D) Spatially contiguous adipose-rich groupings are observed.

**Figure S2.**
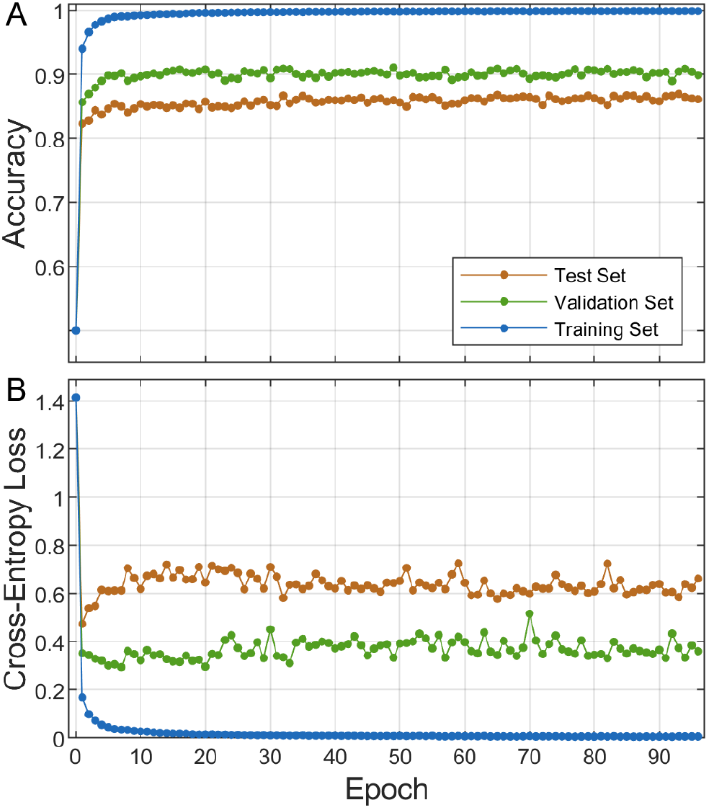
Convergence of an ANN model for a representative RRSS repetition, extended to 96 epochs. A) Accuracy obtained vs. epoch for the training set (blue), a validation set (green), and a test set (orange). B) Cross-Entropy loss vs. epoch for the same data. Note that RRSS rep test set was subdivided randomly by image (16 C, 16 NC in each set) into validation and test sets.

**Figure S3.**
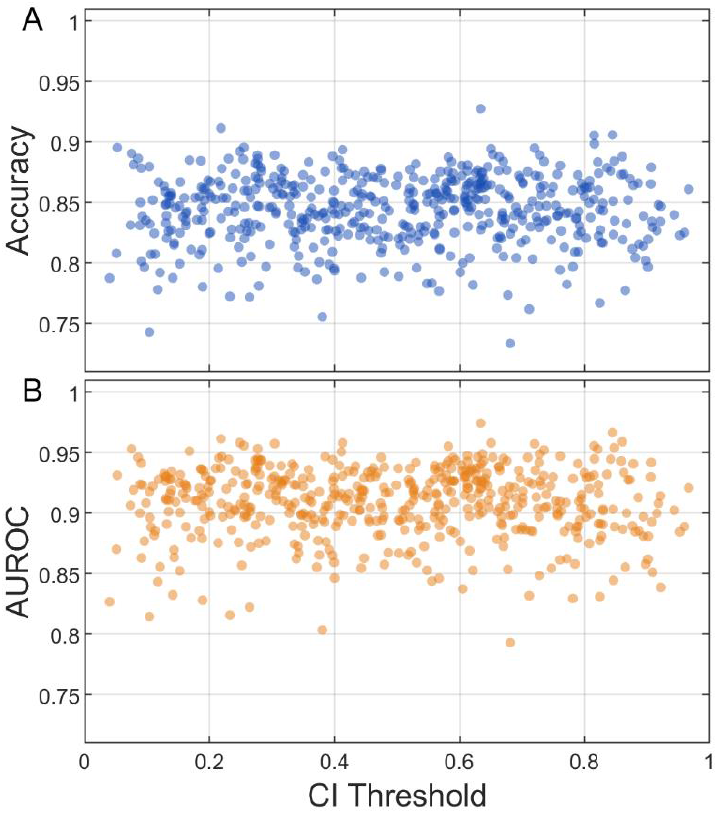
Classification quality behavior vs CI threshold. A) Accuracies vs. corresponding CI thresholds that produced the accuracies. B) AUROCs of reps vs. corresponding CI thresholds.

## Notes

### Competing Interest Statement

The authors have declared no competing interest.

